# The immunosuppression of macrophages underlies the cardioprotective effects of catestatin (CST)

**DOI:** 10.1101/2020.05.12.092254

**Authors:** Wei Ying, Kechun Tang, Ennio Avolio, Jan M. Schilling, Teresa Pasqua, Matthew A. Liu, Hongqiang Cheng, Hong Gao, Jing Zhang, Sumana Mahata, Myung S. Ko, Gautam Bandyopadhyay, Soumita Das, David M. Roth, Debashis Sahoo, Nicholas J.G. Webster, Farah Sheikh, Gourisankar Ghosh, Hemal H. Patel, Pradipta Ghosh, Geert van den Bogaart, Sushil K. Mahata

## Abstract

Hypertension (HTN) is associated with inflammation and excessive production of catecholamines. Hypertensive patients have reduced plasma levels of Catestatin (CST), a bioactive cleavage product of the prohormone Chromogranin A (CgA). In mouse models, HTN symptoms can be reduced by administration of CST, but the role of CST in the regulation of cardiovascular function is unknown. In this study, we generated mice with knockout (KO) of the region of the CgA gene coding for CST (CST-KO) and found that CST-KO mice are not only hypertensive as predicted, but also display left ventricular hypertrophy, have marked macrophage infiltration of the heart and adrenal gland, and have elevated levels of pro-inflammatory cytokines and catecholamines. Intraperitoneal injection with CST reversed these phenotypes, and ischemic pre-conditioning-induced cardioprotection was also abolished in CST-KO mice. Experiments with chlodronate depletion of macrophages and bone-marrow transfer showed that macrophages produce CST and that the anti-hypertensive effects of CST are mediated in part via CST’s immunosuppression of macrophages as a form of feedback inhibition. The data thus implicate CST as a key autocrine attenuator of the cardiac inflammation in HTN by reducing macrophage inflammation.

## Introduction

Hypertension (HTN) is an important risk factor for cardiovascular disease and mortality ^1^. The burden of HTN and the estimated HTN-associated deaths have increased substantially over the past 25 years. The immune system is well recognized for the genesis and progression of HTN ^2, 3^. Compared to healthy individuals, hypertensive or pre-hypertensive patients have elevated levels of pro- and reduced levels of anti-inflammatory cytokines ^4-6^. Inflammatory cytokines can lead to vascular and renal dysfunction and progression of HTN ^7^. Moreover, inflammatory cytokines can increase blood pressure (BP) by increasing the production of catecholamines in the adrenal gland. Specifically, studies in animal models and cultured neuroendocrine cells show that inflammatory cytokines such as interleukin (IL)-1β, interferon (IFN)-α, IL-6 and tumor necrosis factor (TNF)-α can elevate production of norepinephrine (NE), and epinephrine (EPI) ^3, 8, 9^. The dysregulation of the production of catecholamines has been well recognized in HTN ^10, 11^.

Here, we reveal how catecholamine production is attenuated by another secretion product of neuroendocrine cells: the peptide catestatin (CST). CST is a bioactive proteolytical fragment from the pro-hormone Chromogranin A (CgA; hCgA_352-372_) ^12^, which is co-stored and co-released with catecholamines in neuroendocrine cells ^13^. Likely as a consequence of higher catecholamine production ^3^, CgA levels are elevated in humans with essential HTN ^14^ and in rodent genetic models of HTN ^14^. However, unlike CgA, plasma CST levels are diminished not only in essential HTN ^14, 15^, but also in the normotensive offspring of patients with HTN ^15^, suggesting dysregulation in the processing of CgA to CST ^14^. Moreover, HTN-associated single nucleotide polymorphisms within the CST segment of CgA have been identified ^16-18^.

Animal experiments also indicate a role for CST in HTN: both CgA heterozygote and complete knockout (KO; CgA-KO) mice are hypertensive, and treatment with CST normalizes the BP and plasma catecholamine levels ^19^. In mouse models of diabetes ^20^, colitis ^21^ and atherosclerosis ^22^, CST exerts anti-inflammatory effects by inhibiting the activation of macrophages and shifting their differentiation to more anti-inflammatory phenotypes ^23^. Therefore, we hypothesized that CST exerts its cardioprotective role by skewing macrophages to more anti-inflammatory phenotypes, thereby resulting in lower catecholamine production.

To directly discern the role of CST in the regulation of the cardiovascular system, we generated CST-KO mice, which lack only the CST-coding region of the *Chga* gene. As predicted, CST-KO mice displayed a hypertensive, hyperadrenergic, and inflammatory phenotype which was rescued by exogenous addition of CST. Macrophage depletion by chlodronate (CDN) liposomes and from bone-marrow transfer (BMT) between CST-KO and wild-type (WT) mice revealed that macrophages produce CST and are responsible for the anti-inflammatory/anti-HTN effects of CST. CST might be a novel target for the treatment and prevention of HTN.

## METHODS

Full Materials and Methods are available in the Data Supplement. Data and protocols are also available upon reasonable request from the corresponding author.

### Mice

We used male WT and CST-KO (20-24 weeks old) in C57BL/6 background unless indicated otherwise. Since CgA is especially overexpressed in male patients with hypertension ^24^, we used only male mice in this study. Mice were kept in a 12 hr dark/light cycle and fed a normal chow diet (NCD: 13.5% calorie from fat; LabDiet 5001, TX). Animals were age-matched, and randomly assigned for each experiment. Control and experimental groups were blinded. Power calculations were conducted to determine the number of mice required for each experiment. For rescue experiments, mice were injected intraperitoneally with CST (2 µg/g body weight) at 9:00 AM for 2-4 weeks before harvesting tissues. All studies with mice were approved by the UCSD and Veteran Affairs San Diego Institutional Animal Care and Use Committees and conform to relevant National Institutes of Health guidelines.

### Statistics

Statistics were performed with PRISM 8 (version 8.4.3) software (San Diego, CA). Data were analyzed using unpaired two-tailed Student’s *t*-test for comparison of two groups or one-way or two-way analysis of variance (ANOVA) for more than two groups followed by Tukey’s *post hoc* test if appropriate. All data are presented as mean ± SEM. Significance was assumed when p<0.05.

## RESULTS

### Generation and validation of CST-KO mice

The CST coding region (mCgA_364-384_; 63 bp) was removed from Exon VII of the *Chga* gene (Figure 1A&B). Using a mouse monoclonal antibody (5A8), we detected full-length CgA (∼70 kDa) in WT mice and a proteoglycan form of CgA in CST-KO mice in adrenal gland lysates (Figure 1C), indicating the presence of CgA in CST-KO mice. Blots using a polyclonal antibody directed against the C-terminal domain of CST (CT-CST) showed a proteolytically processed CgA (∼46 kDa) corresponding to mCgA_1-385_ in WT littermates, but not in CST-KO mice (Figure 1C). Because this antibody detects synthetic CST (positive control for antibody specificity), we conclude that CST-KO mice indeed lack CST. Adrenal CgA content was comparable in WT and CST-KO mice (Figure 1D). CST was not detectable in CST-KO mice (Figure 1D).

**Fig 1.**
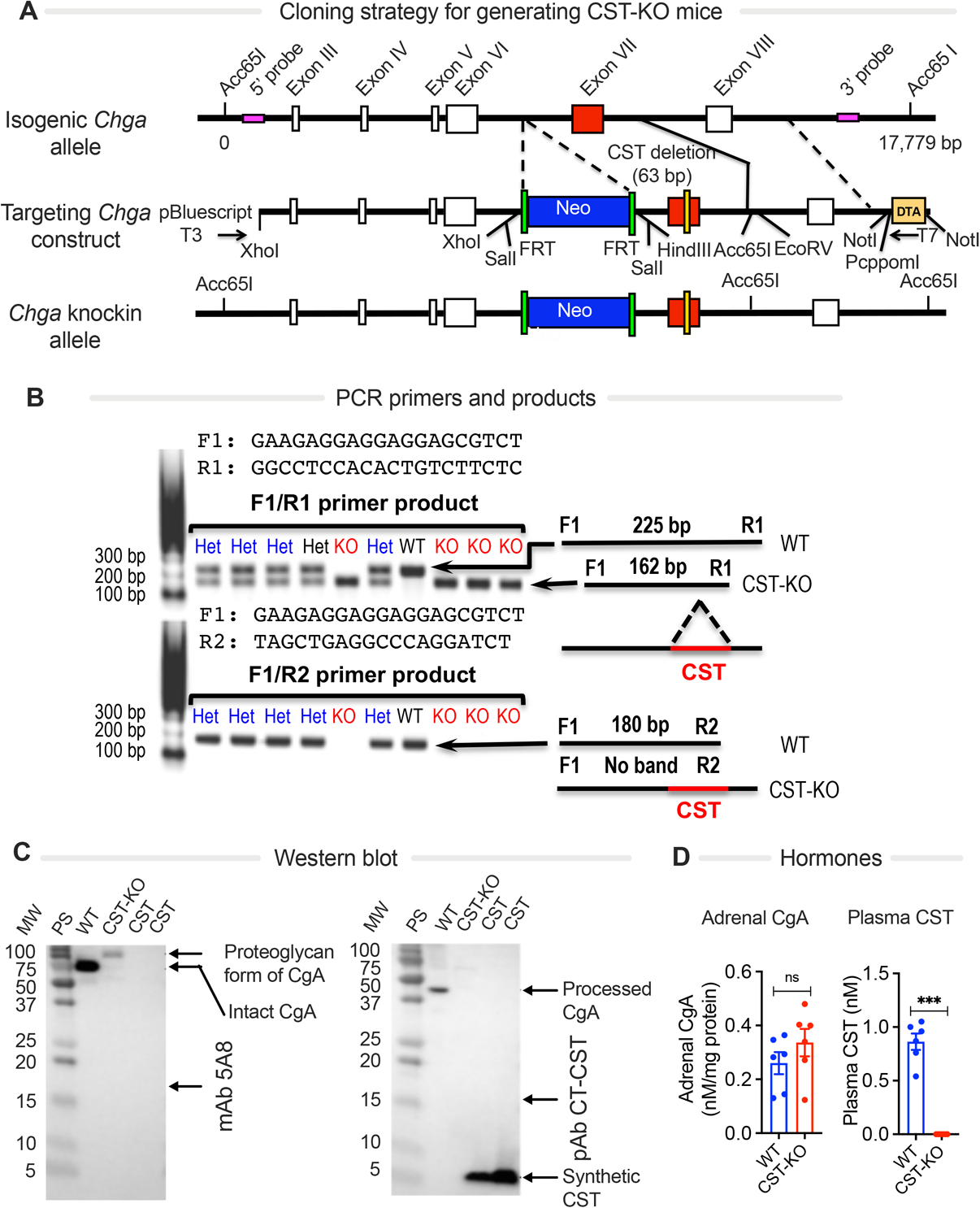
Generation of CST-KO mice. **(A)** Schematic diagram showing the cloning strategy for generating CST-KO mice. DTA, diphtheria toxin; FRT, *Flp* recognition target. **(B)** Screening of CST-KO mice by PCR. Primer set 1 flanks the CST domain; expected PCR products: 162 bp, CST-KO; 225 bp, WT mice. Reverse primer 2 binds within CST-coding region; no band, CST-KO, 180 bp WT mice. **(C)** Western blots showing the presence of CgA and CST in WT and CST-KO mice using monoclonal antibody (mAb) 5A8 that does not recognize CST, and rabbit polyclonal antibody directed against the C-terminus of CST (pAb CT-CST), which does not recognize CgA beyond CST domain. A truncated CgA (CgA_1-384_) was present in WT mice but not in CST-KO mice, confirming the deletion of CST domain. **(D)** Adrenal CgA content (n=12) and plasma CST levels (n=6). Unpaired two-tailed *t*-test: ns, not significant; ***p<0.001.

### CST-KO mice are hypertensive

Consistent with the anti-HTN functions of CST ^19, 25, 26^, we found that the CST-KO mice were hypertensive and displayed diurnal increases in both systolic and mean arterial BP (Figure 2A & S1). The high BP in CST-KO mice was rescued by intraperitoneal injection of exogenous CST (2 µg/g body weight for 15 days), whereas CST did not affect normotensive BP in WT mice (Figure 2B). In WT mice, the plasma CST level was 0.86 nM, which increased to 1.72 nM 24 hrs after administration of CST (Figure 2C). In CST-KO mice, plasma CST was 1.17 nM after 24 hr of CST supplementation, indicating that CST supplementation provided a near physiological concentration of CST.

**Fig 2.**
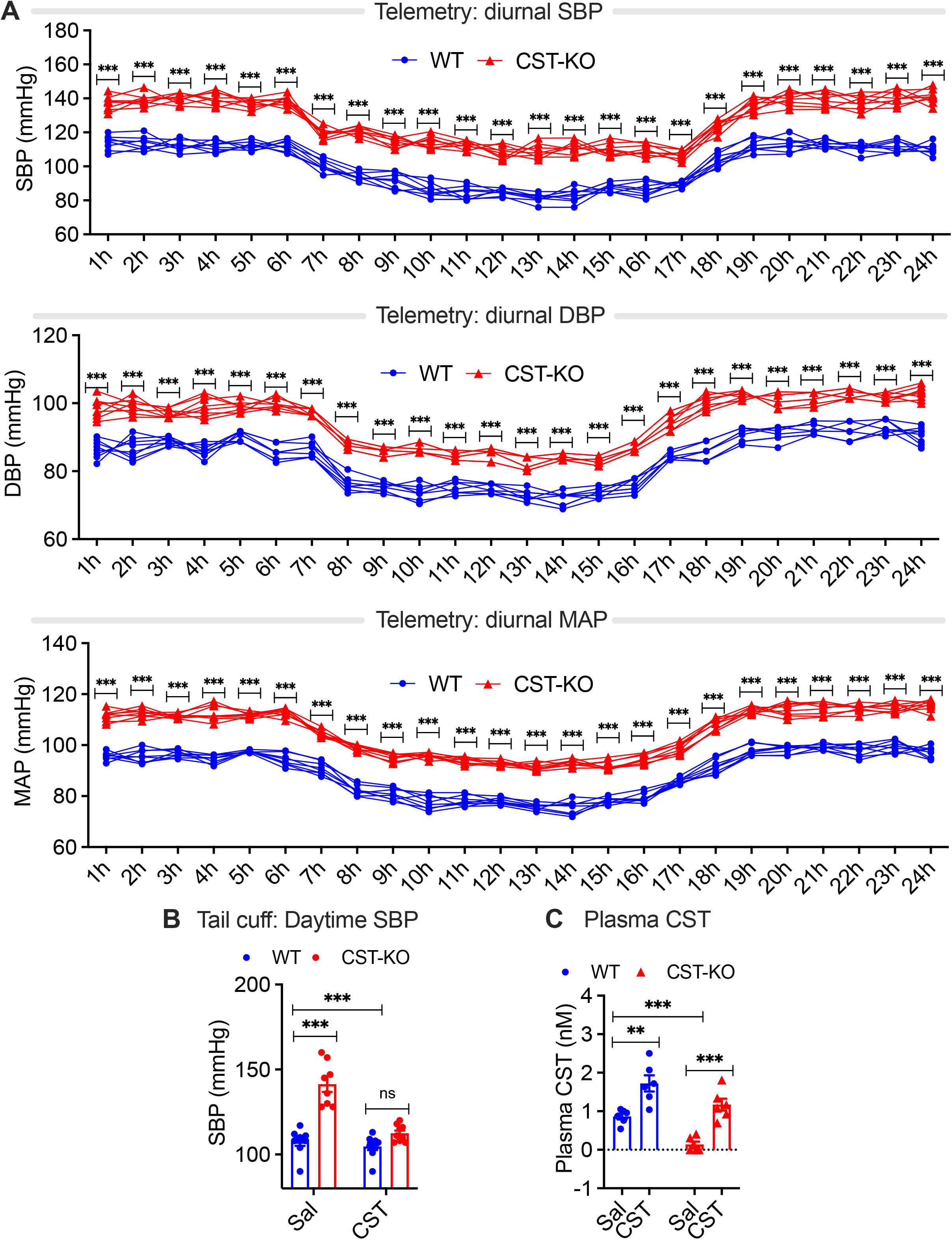
Hypertension in CST-KO mice. **(A)** Diurnal systolic (SBP), diastolic (DBP) and mean arterial (MAP) blood pressure by telemetry in wild-type (WT) and CST-KO mice (n=8). **(B)** Daytime SBP by tail-cuff (n=9) and **(C)** plasma CST levels (n=6) of WT and CST-KO mice treated with CST or saline (Sal). Two-way ANOVA: ns, not significant; **p<0.01; ***p<0.001.

### Ischemic pre-conditioning-induced cardioprotection is impaired in CST-KO mice

Since CST promotes cardioprotection in rats ^27^, we tested whether pre-conditioning-induced cardioprotection is affected in CST-KO mice. We subjected WT and CST-KO hearts to ischemia/reperfusion (IR) followed by ischemic preconditioning (IPC). IPC significantly increased the post-ischemic left ventricular developed pressure (LVDP) and lowered the left ventricular end diastolic pressure (LVEDP) in WT hearts compared to CST-KO hearts and their respective IR controls (Figure S2A). Furthermore, neither LVDP nor LVEDP was significantly modified in IPC-treated CST-KO hearts compared to the respective IR controls. In WT mice, but not in CST-KO mice, IPC also improved recoveries of both the maximum and minimum rates of pressure development in the LV (dP/dt_max_ and dP/dt_min_) compared to the respective IR controls (Figure S2B). These data show that whereas IPC conferred protection against IR damage in WT mice, CST-KO mice could not be preconditioned.

### CST-KO mice have increased inflammation in heart and circulation

CST is an anti-inflammatory peptide ^23 28^, raising the possibility that CST might regulate cardiovascular function via the immune system. Indeed, in plasma of CST-KO mice, we found increased levels of proinflammatory cytokines TNF-α, IFN-γ, C-C motif chemokine ligand (CCL)-2 and -3, and C-X-C motif chemokine ligand (CXCL)-1 (Figure 3A). By contrast, the anti-inflammatory cytokine IL-10 was decreased in CST-KO mice. Intraperitoneal injection with exogenous CST in CST-KO mice reversed this phenotype: it decreased the levels of most proinflammatory cytokines and increased anti-inflammatory cytokines in plasma of both WT and CST-KO mice (Figure 3A). RT-PCR also revealed inflammation in the heart of CST-KO mice: the expression of anti-inflammatory genes *IL10, IL4, Mrc1, Arg1, Clec7a* and *Clec10a* was reduced, whereas the pro-inflammatory genes *Tnfa, Ifng, Emr1, Itgam, Itgax, Nos2a, IL12b CcL2*, and *CxcL1* were upregulated (Figure 3B&C). LV protein levels of the proinflammatory cytokines TNF-α, IFN-γ, CCL-2, CCL-3, CXCL-1, and IL-6 were also elevated in CST-KO mice (Figure 3D). These phenotypes were also reversable by intraperitoneal injection of CST. We also observed increased phosphorylation (Ser177/181) of IKK-β, a component of the inflammatory NF-κB signaling pathway (Figure 4A). These findings show that the immune system of CST-KO mice is skewed towards inflammation.

**Fig 3.**
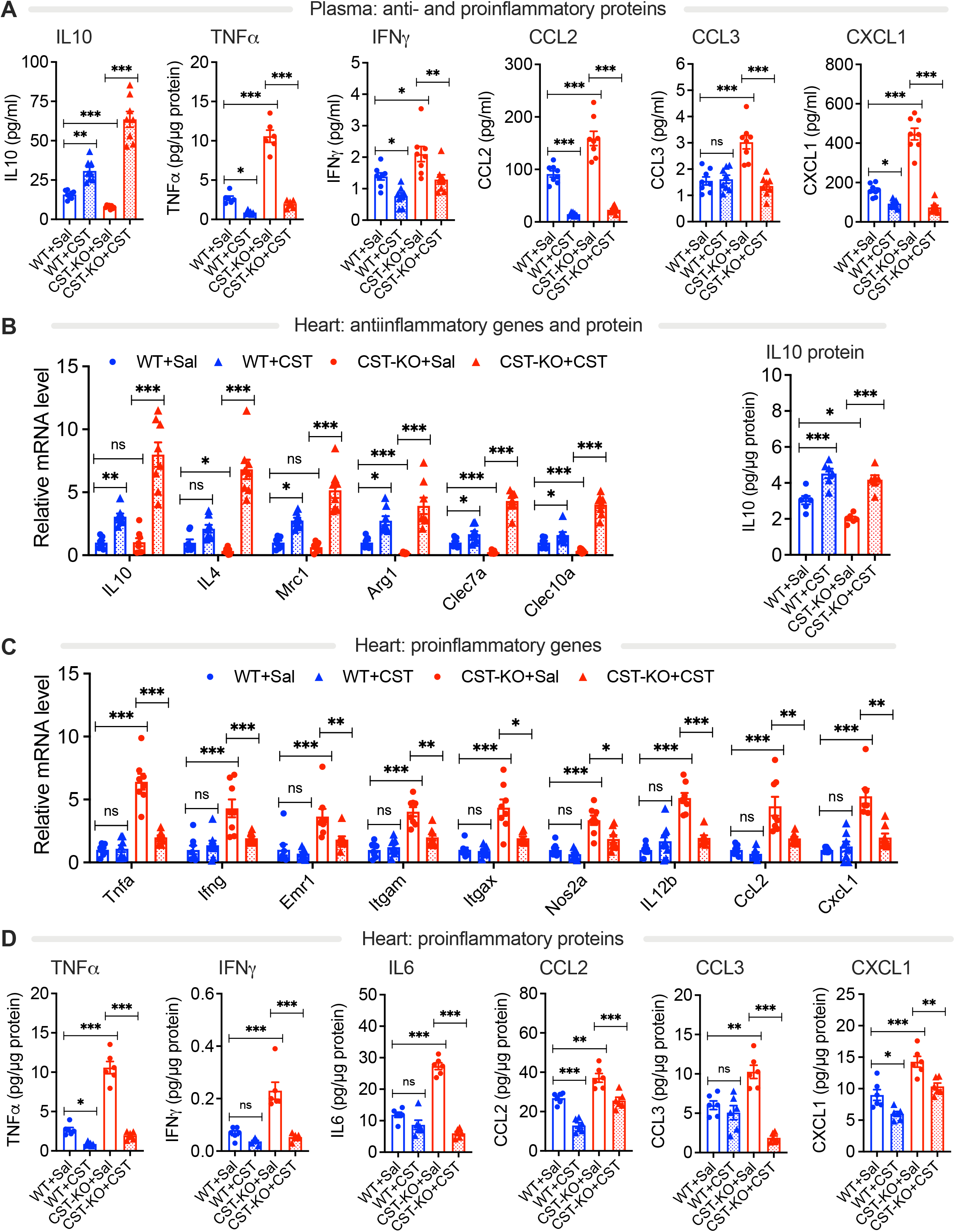
CST-KO mice display systemic and cardiac inflammation, which can be reversed with exogenous CST. **(A)** Plasma cytokines in WT and CST-KO mice (n=8). Sal: saline. **(B&C)** RT-qPCR data showing steady-state mRNA levels of anti-**(B)** and pro-inflammatory **(C)** genes in left ventricle (n=8). **(B&D)** Protein levels of IL-10 **(B)** and pro-inflammatory cytokines **(D)**. One-way ANOVA: ns, not significant; *p<0.05; **p<0.01; ***p<0.001.

**Fig 4.**
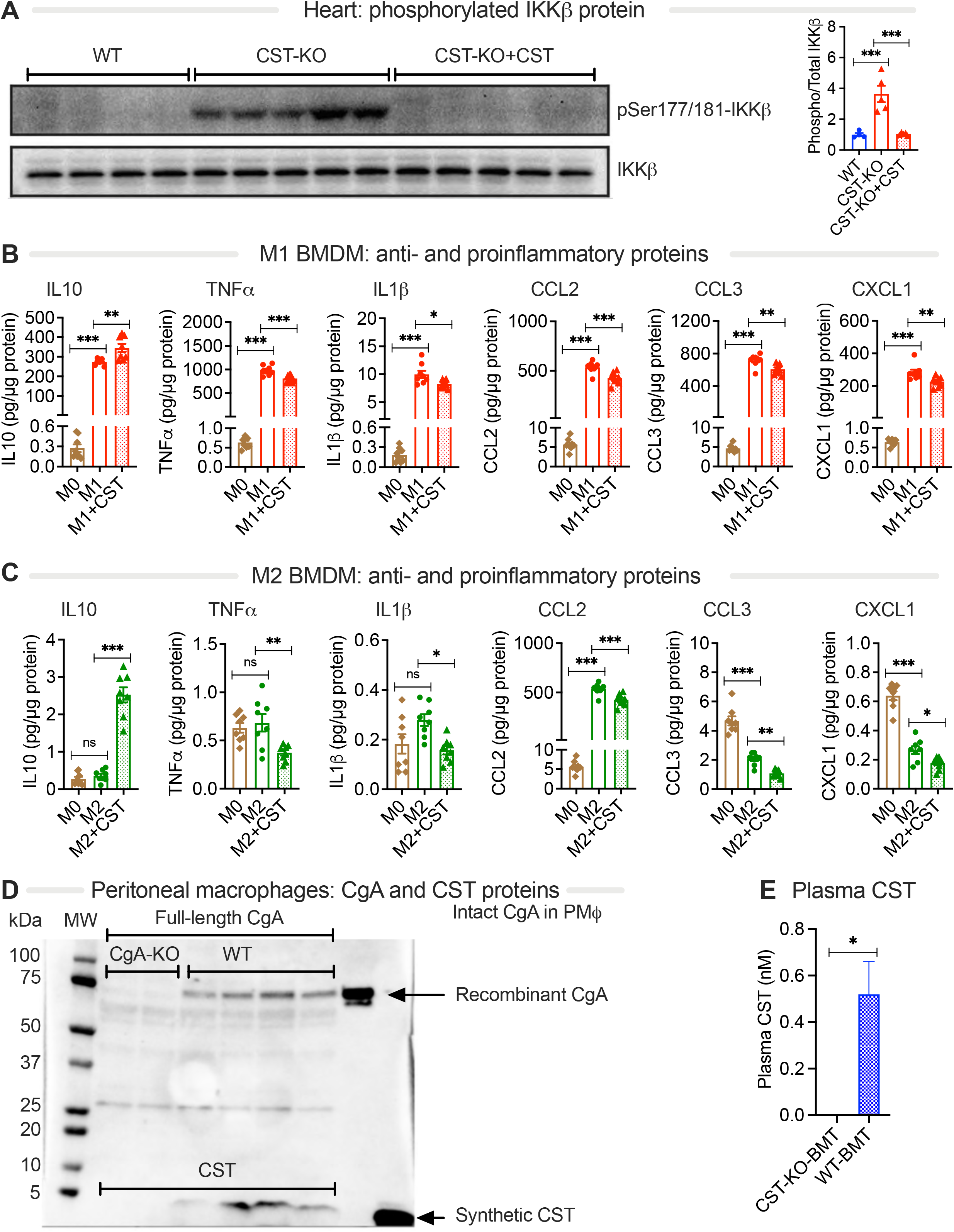
CST exerts anti-inflammatory effects in isolated macrophages and macrophages are major producers of CST. **(A)** Western blot analysis of phosphorylated (Ser177/181) and total IKK2 in heart of 4 WT mice, 5 CST-KO mouse and 5 CST-KO mice with intraperitoneal injection of CST. **(B&C)** Protein levels of cytokines (n=8) in supernatant of bone-marrow derived macrophages (M0-BMDMs) differentiated to proinflammatory M1-type **(B)** and anti-inflammatory M2-type phenotypes **(C)**. Cells were treated with 100 nM CST. **(D)** Western blots showing the presence of CgA and CST in peritoneal macrophages (n=4). **(E)** Plasma CST levels in CST-KO mice which received BMT from CST-KO or WT mice (n=6). Panel A-C: one-way ANOVA; panel E: *t*-test; ns, not significant; *p<0.05; **p<0.01; ***p<0.001.

### CST reduces pro-inflammatory macrophages *in vitro*

In the next set of experiments, we addressed whether CST would directly shift macrophages to more anti-inflammatory responses *in vitro*. Macrophages were derived from bone-marrow of WT mice (BMDM) and differentiated to an either a pro-inflammatory M1-like phenotype or an anti-inflammatory M2-like phenotype ^29^. Culturing these macrophages for 24 hr with 100 nM CST resulted in a small, but significant, reduction of the production of pro-inflammatory cytokines TNF-α, CCL-2, CCL-3, CXCL-1 and IL-1β (Figure 4B&C). In contrast, the levels of anti-inflammatory IL-10 were increased in the macrophages.

Since macrophages are a secretory cell type, we also addressed whether macrophages produce CgA and CST. Indeed, Western blotting analysis revealed the presence of both CgA and CST in cultured peritoneal macrophages (Figure 4D) ^29^. To assess the physiological relevance of CST-production by macrophages, we performed BMT experiments in which we irradiated CST-KO mice and then cross-transplanted the marrow from WT mice. We analyzed plasma CST of these mice and found that WT bone-marrow recipient CST-KO mice, but not CST-KO bone-marrow recipients, had near physiologic levels of plasma CST (0.52 nM) (Figure 4E). Thus, macrophages (and possibly other bone-marrow derived cell types) are major producers of CST in circulation.

### Macrophages are key effector cells responsible for the anti-inflammatory actions of CST

TEM studies revealed abundant infiltration of macrophages and fibrosis in the intercellular spaces between chromaffin cells in the adrenal medulla of CST-KO mice (Figure S3 and S4). Also, marked cardiac fibrosis and an increased presence of recruited monocytes were observed in the heart of saline-treated CST-KO mice, as shown by TEM and flow cytometry (Figure 5A, S3, S5 and S6A). In both the adrenal gland and heart of CST-KO mice, CST supplementation reduced the abundance of macrophage infiltrates (Figure S3). Flow cytometry analysis showed a decrease of the fraction of F4/80^-^Ly6C^+^ monocytes compared to total CD11b^+^ cells in CST-supplemented CST-KO heart (Figure 5A and S6B).

**Fig 5.**
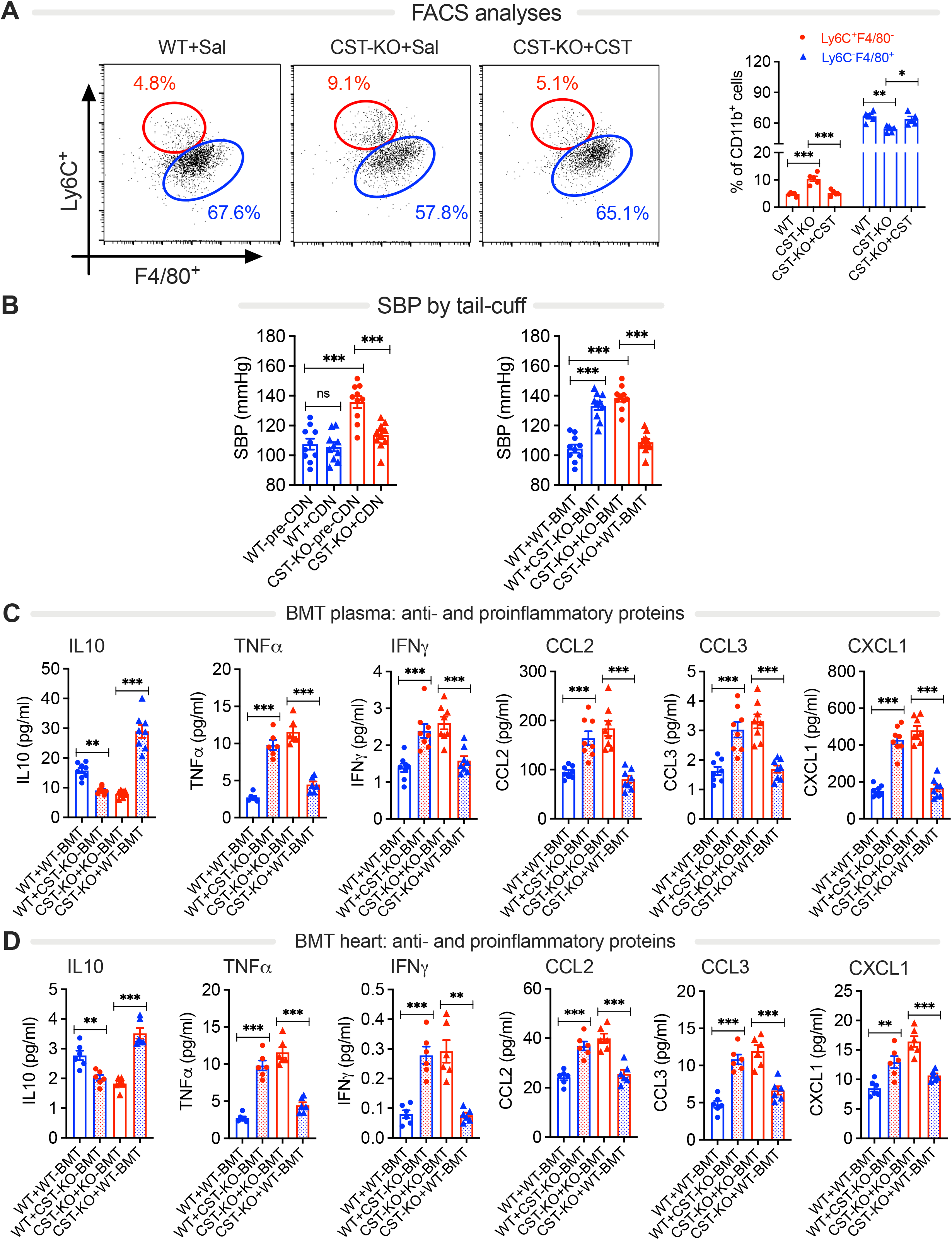
Macrophages mediate cardioprotective effect of CST. **(A)** Flow cytometry data showing CD45^+^F4/80^+^CD11b^+^Ly6c^-^ macrophages and CD45^+^F4/80^-^CD11b^+^Ly6c^+^ recruited monocytes in CST-KO heart (n=3). Sal: saline control. **(B)** Systolic blood pressure (SBP) after depletion of macrophages by CDN (n=8) and bone-marrow transfer (BMT) into irradiated mice (n=8). Levels of cytokines in plasma **(C)** and heart **(D)** of mice with BMT (n=8). One-way ANOVA: ns, not significant; *p<0.05; **p<0.01; ***p<0.001.

We assessed the functional role of the infiltrated macrophages in the heart and adrenal gland of CST-KO mice using two independent approaches. First, we depleted macrophages by CDN liposomes (Figure S6B), which not only depleted macrophages in heart and adrenal gland (Figure S3), but also reversed the hypertensive phenotype of CST-KO mice (Figure 5B). Second, we carried out BMT assays in which we irradiated both WT and CST-KO mice and then cross-transplanted their marrows: bone-marrow from CST-KO mice was transplanted into WT mice and *vice versa*. Both the inflammatory and hypertensive phenotypes were transferred by BMT: while CST-KO bone-marrow recipient WT mice showed increased BP; elevated levels in plasma and heart of TNF-α, IFN-γ, CCL-2, CCL-3, and CXCL-1; and reduced levels of IL-10, WT bone-marrow recipient CST-KO mice showed the opposite phenotypes (Figure 5B-D). Since WT bone-marrow recipient CST-KO mice had near physiologic levels of plasma CST (Figure 4E), we conclude that macrophages and other immune cells are not only key effectors of the anti-hypertensive actions of CST but are also main producers of CST themselves.

### Heightened sympathetic stimulation and hypersecretion of catecholamines in adrenal gland of CST-KO mice

Prior studies have shown that pro-inflammatory cytokines increase catecholamine production and secretion ^3, 8, 9^. In line with this, we found that compared to WT littermates, CST-KO mice have elevated levels of both adrenal and plasma catecholamine levels (Figure 6A). This phenotype was also transferable by BMT: CST-KO bone-marrow recipient WT mice showed increased levels of NE and EPI in the adrenal medulla and plasma, whereas WT bone-marrow recipient CST-KO mice showed the opposite phenotype (Figure 6A).

**Fig 6.**
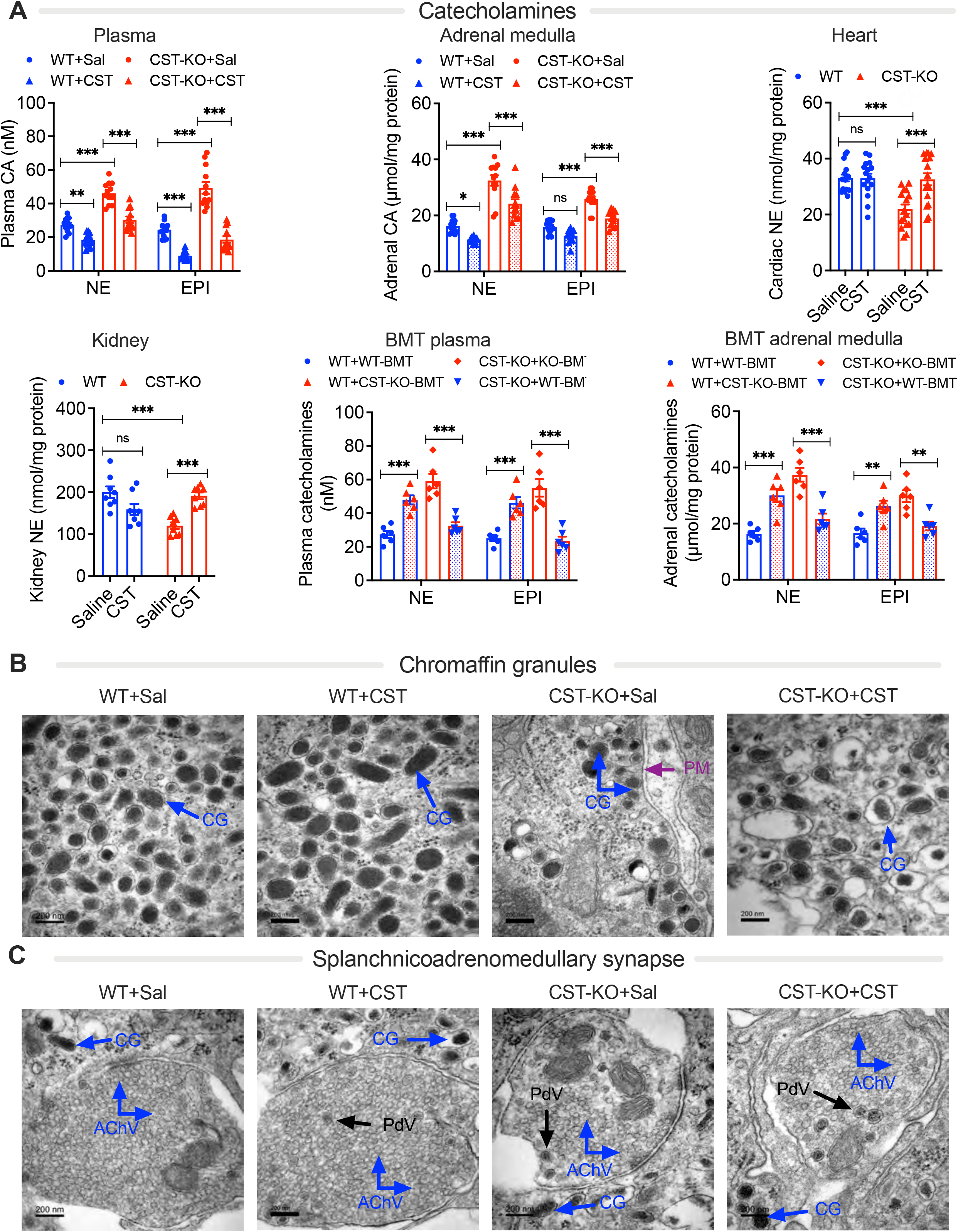
Increased catecholamine secretion in CST-KO mice. **(A)** Plasma, adrenal, heart and kidney norepinephrine (NE) and epinephrine (EPI) levels in WT and CST-KO mice. Mice were treated with saline (Sal) or CST or underwent bone-marrow transfer (BMT) into irradiated mice (n=8-12). **(B)** TEM micrographs of chromaffin granules (CG) in the adrenal medulla of CST-KO mice. PM, plasma membrane. **(C)** TEM micrographs of splanchnic-adrenomedullary synapse with acetylcholine vesicles (AChV) and dense core peptidergic vesicles (PdV). CC, chromaffin cell; CG, chromaffin granule. Two-way ANOVA: ns, not significant; *p<0.05; **p<0.01; ***p<0.001.

Since heightened sympathetic nerve traffic has been documented in hypertensive patients ^30^, we measured NE in the LV and kidney of WT and CST-KO mice. In contrast to the adrenal medulla and plasma, we observed reduced levels of NE in the heart and kidney of CST-KO mice (Figure 6A). Decreased NE in CST-KO mice indicates increased cardiac and renal spillover of NE, which is common in hypertensive and heart failure patients ^31-34^. The CST-KO adrenal medulla exhibited abundant docked chromaffin granules and decreased acetylcholine-containing vesicles at the sympatho-adreno-medullary synapse (Figure 6B-C and S7), implicating heightened sympathetic nerve activity leading to hypersecretion of catecholamines ^35, 36^. Supplementation of CST-KO mice with CST reversed this phenotype (Figure 6B-C and S7) and led to a concomitant decrease in both plasma and adrenal catecholamines (Figure 6A). Since CST acts as an endogenous inhibitor of nicotine-induced catecholamine secretion ^37^, we reasoned that ganglionic blockade would reverse elevated BP in CST mice. Indeed, the elevated SBP, DBP, and MAP in CST-KO mice were reversed by the long-acting nicotinic acetylcholine receptor antagonist chlorisondamine (Figure S8A-C). This compound also decreased the heart rate (HR) (Fig. S8D) as reported by others ^38^.

To test whether lack of CST affected heart structure and function, we undertook gravimetry and echocardiographic ultrasound imaging. CST-KO mice showed increased heart weights and sizes compared to WT mice (Figure S9A). Although CST-KO mice maintained a similar level of LV function (fractional shortening) to age-matched WT mice, there were significant abnormalities in LV remodeling as evidenced by the significant increase in LV posterior wall thickness (LVPWd), which has been associated with high BP ^39, 40^, and a trend towards an increase in interventricular septum wall thickness (IVSd; p=0.07) (Figure S9B). HR, LV internal diameter during systole, and LV internal diameter during diastole were comparable between WT and CST-KO mice (Figure S9B).

## DISCUSSION

### Inflammation and hypertension

Inflammation is well understood to contribute to the development of hypertension by inducing vascular damage, renal damage, and/or abnormal central neural regulation ^41-43^. This study showed that inflammation and the development of HTN are counteracted by the anti-inflammatory hormone CST through its feedback inhibition/regulation of macrophages and the anti-inflammatory actions of CST are likely needed to maintain the tolerance of the immune system. Previous studies showing low levels of CST in hypertensive subjects ^15^ and normalization of BP in CgA-KO mice by CST ^19^ as well as decreasing BP in spontaneously hypertensive rats ^25^ indicate that CST is sufficient to reverse HTN. The findings from this study that CST-KO mice are hypertensive with a skewed immune system towards inflammation, and that these phenotypes can be rescued by exogenous administration of recombinant CST, add to this and demonstrate that CST is not only sufficient but also necessary for regulating HTN. Since the inflammation and BP can be reduced by administration of exogenous CST, CST might be a therapeutic target for the treatment of HTN.

Suppression of the immune system attenuates the development of HTN when induced by Angiotensin II or deoxycorticosterone acetate-salt, while dysregulation of it causes sensitization to these hypertensive challenges ^44^. Surprisingly, CST-KO mice already show elevated BP in absence of an additional challenge, raising the question of other mechanisms in addition to the immune activation are involved in the HTN phenotype in these mice. However, since bone-marrow transplant from CST-KO to WT mice already suffices to elicit HTN, and these mice only harbor CST lacking immune cells while the autonomic system is normal, it might be that an abnormal activation of the immune system triggers HTN.

### Neurohumoral regulation of BP

We found a neuro-adrenergic overdrive-induced HTN in CST-KO mice. Existing literature reveals heightened sympathetic nerve activity in hypertensive subjects ^45, 46^. In addition, hypertensive patients with metabolic risk factors, such as obesity, metabolic syndrome, or diabetes mellitus, also exhibit sympathetic overdrive ^47-49^. Like humans, spontaneously hypertensive rats show reduced cardiac parasympathetic nerve activity, elevated sympathetic nerve activity and increased NE release ^50^.

### Immunoendocrine regulation of BP

The augmented sympathetic nerve activity in HTN is known to activate both myeloid cells and T cells ^2^, and circulating concentrations of pro-inflammatory cytokines are increased in primary HTN ^51^. T-lymphocytes are critical for Angiotensin II and deoxycorticosterone acetate-salt-induced hypertension ^52, 53^. Intracerebroventricular administration of IL-6 increases splenic sympathetic nerve activity ^54^, while central administration of IL-1β increases adrenal, splenic and renal sympathetic nerve activity ^55^. Injection of TNF-α into central sympathetic nuclei, such as the paraventricular nucleus increases sympathetic nerve activity, BP and heart rate in rats ^56^.

To our knowledge, the present study is the first to demonstrate increased infiltration of macrophages in the adrenal medulla concomitant with increased secretion of catecholamines and the consequent development of HTN in CST-KO mice, which were normalized after CST supplementation. These findings imply that CST regulates the BP through a novel immunoendocrine regulation of catecholamine secretion via macrophages.

What causes the elevated BP in CST-KO mice? Since nicotine-induced increases in BP are associated with an increase in plasma catecholamine secretion ^57^, it is consistent that lack of CST (an endogenous inhibitor of nicotine-evoked catecholamine secretion ^37^) in CST-KO mice resulted in increased plasma catecholamines with consequent increase in BP. We have confirmed this by using chlorisondamine, which decreased BP in CST-KO mice as has been reported in rats ^38^. However, in CST-KO mice, the HR was not increased, and fractional shortening was unaltered compared to WT, indicating that the increased BP is not driven by elevated cardiac output. Further studies are required to explain what causes hypertension in CST-KO mice. It might be that the LV hypertrophy in the CST-KO mice, and possibly also the increased posterior wall thickness associated with high BP ^39, 40^, is a secondary effect of the BP elevation. It seems therefore possible that the elevated BP is caused by increased vascular resistance, due to vasoconstriction and/or increased arterial stiffness ^58^. It is possible that increased cardiac and renal spillover of NE also contribute to the development of BP in CST-KO mice ^31-34^.

### Perspectives

This study provides a key mechanism as to how CST regulates inflammation. Cardiac macrophages are critical for myocardial homeostasis ^59, 60^. We found an abundance of infiltrated macrophages in the heart and adrenal gland of CST-KO mice. Using *in vitro* experiments with cultured BMDMs and two parallel approaches (CDN macrophage depletion and BMT) to decrease macrophage activity *in vivo*, we found that macrophages are key effector cells for the anti-hypertensive actions of CST. We also found that bone-marrow originated cells, possibly macrophages, are the main source of circulating CST.

We therefore propose that the macrophages (and chromaffin cells) produce CST, which reduces inflammation by autocrine/feedback inhibition fashion. These anti-inflammatory actions underlie the anti-hypertensive effects of CST, since without CST, macrophages are more reactive, infiltrate the heart, and alter the ultrastructure, physiological makeup, and molecular makeup of the myocardium. Additionally, the data implicate CST as a key mediator of the observed crosstalk between systemic and cardiac inflammation in HTN, which hence plays a central role in cardiovascular homeostasis by regulating the immunoendocrine axis.

## Supporting information

Supplemental methods_Figures_BioRxiv_022521

## ACKNOWLEDGEMENTS

None

## NOVELTY AND SIGNIFICANCE

### What is new?

- Mice that lack the peptide hormone catestatin (CST) are hypertensive.
- CST skews macrophages to an anti-inflammatory phenotype.
- CST reduces inflammation in heart and adrenal gland.

### What is relevant?

- Hypertension is associated with inflammation.
- Hypertensive patients have reduced plasma levels of CST.
- In mouse models, hypertension can be reduced by exogenous CST.

### Summary

- The anti-hypertensive effects of CST are mediated via CST’s immunosuppression of macrophages.
- CST is a key autocrine attenuator of the cardiac inflammation in hypertension.

## ABBREVIATIONS

BMDM: bone marrow-derived macrophage
BP: blood pressure
BMT: bone marrow transfer
CCL: C-C motif chemokine ligand
CXC: C-X-C motif chemokine ligand
CDN: chlodronate
CgA: chromogranin A
CST: catestatin
EPI: epinephrine
HTN: hypertension
IFN: interferon
IL: interleukin
IPC: ischemic preconditioning
IR: ischemia/reperfusion
KO: knockout
LV: left ventricular
LVDP: LV developed pressure
LVEDP: LV end diastolic pressure
LVPWd: LV posterior wall thickness
MAP: mean arterial pressure
IVSd: interventricular septum wall thickness
NE: norepinephrine
SBP: systolic blood pressure
TEM: transmission electron microscopy
TNF: tumor necrosis factor
WT: wild-type

