## Supplemental methods_Figures_BioRxiv_022521 for "The immunosuppression of macrophages underlies the cardioprotective effects of catestatin (CST)"

### SUPPLEMENTARY METHODS

**Protein analysis by immunoblotting.** Left ventricles were homogenized in a buffer containing phosphatase and protease inhibitors, as previously described [1]. Peritoneal macrophages (PMϕ) were prepared and cultured as previously described. Ventricular homogenates and PMϕ lysates were subjected to SDS-PAGE and immunoblotted with Mouse monoclonal mAb 5A8 ($hCgA_{R47-L57;}$ 1:500; generously provided by Angelo Corti, Milan, Italy) antibody, rabbit polyclonal C-terminal CST ($hCgA_{P368-R373}$; 1:500; generously provided by Angelo Corti, Milan, Italy) [2], rabbit anti-phospho-IKKa/b (Ser176/180; 1:5000; Cell Signaling, USA) or rabbit anti-IKK2 (1:5000; BioBharati LifeScience, India).

**Masson's trichrome staining.** Masson's Trichrome staining technique for the detection of collagen fibers was performed by the Microscopy & Histology core (La Jolla Institute of Allergy and Immunology, La Jolla, CA). Formalin (10%)-fixed heart tissues were embedded in paraffin. Sections (5 µm) were stained with Weigert's hematoxylin (stains nuclei), Biebrich scarlet-acid fuchsin (stains cytoplasm and muscle), and aniline blue (stains collagen). The collagen fibers stained blue and the nuclei stained black, with a red background.

**Transthoracic echocardiography.** Echocardiography was performed (Cardiovascular Physiology Core, UCSD) as previously described [3]. Briefly, adult age-matched WT and CST-KO mice were anesthetized with 5% isoflurane and maintained with 0.5% isoflurane during the entire procedure. The anterior chest was shaved, and small-needle echo leads were entered through the skin on the right and left upper extremities and the left leg. A Vevo 2100 ultrasound system (FUJIFILM VisualSonics Inc.) with a linear transducer with 32-56 MHz range (MS550S) was used to capture the images and videos.

**Telemetric continuous intraarterial measurement of BP**. Diurnal systolic blood pressure (SBP), diastolic blood pressure (DBP), and heart rate (HR) were measured by telemetry using the Data Sciences International (DSI, St. Paul, MN, USA) PhysioTel telemetry system (Cardiovascular Physiology Core, UCSD) as described previously [4, 5].

Telemetric transmitter (TA11PA-C20, DSI) was implanted in the left carotid artery in mice anesthesized with isoflurane (5% for induction and 2% for maintenance). Telemetry signals were received by an antenna below the cage that relayed the data to a signal processor (DataQuest A.R.T. Gold Version 2.3; DSI) connected to a Compaq desktop personal computer (Hewlett-Packard). BP was recorded after 10 days of implantation surgery when the diurnal pattern of BP was normalized in these conscious mice fitted with DSI transmitters.

**Non-invasive tail-cuff measurement of BP**. The SBP was also measured indirectly using tail cuff plethysmography in rat tail-cuff blood pressure system (MRBP; IITC Life Sciences Inc. Woodland Hills, CA). Prior to any measurements, the plexiglass restrainers were warmed in 34°C warming chambers for 20 minutes. The mice were then loaded into their restrainers and allowed to incubate in the same warming chambers for 10-15 min. The tails were placed inside inflatable cuffs with a photoelectric sensor that measured tail pulses. The SBP was measured over 6 separate days with an average of at least two well-defined BP cones detected by the tail cuff plethysmograph per day.

**Determination of plasma CgA and CST.** Commercial mouse EIA kits were used to determine CgA from two adrenal glands (CUSABIO Technology LLC, Houston, TX) and plasma (200 µl) CST (RayBiotech Life, Peachtree Corners, GA) and followed according to the manufacturer's protocol.

**Culturing of BMDMs.** Bone marrow-derived macrophages (BMDMs) were prepared as previously described [6]. Briefly, bone marrow cells were isolated from 6-8 weeks old WT mice, followed by erythrocyte lysis with ammonium chloride, and then seeded in 24-well plates at a concentration of $1x10^6$ cells/ml. Cells were differentiated to macrophages with culture mediums containing monocyte-colony stimulating factor (M-CSF; 10 ng/ml). After 7 days differentiation, naïve (M0) BMDMs were treated with IL4 (20 ng/ml) or LPS (100 ng/ml) with or without CST (100 nM) for 24 hours to stimulate M1 or M2 polarization, respectively.

**Measurement of cytokines.** Plasma cytokines or supernatant of cultured BMDMs (20 µl) were measured using U-PLEX mouse cytokine assay kit (Meso Scale Diagnostics, Rockville, MD) via the manufacturer's protocol.

**Electron microscopy and morphometric analysis.** To displace blood and wash off tissues before fixation, mice were cannulated through the apex of the heart and perfused with a calcium and magnesium free buffer composed of DPBS (Life Technologies Inc), 10 mM HEPES, 0. 2 mM EGTA, 0.2% BSA, 5 mM glucose and KCl concentration adjusted to final 9.46 mM (to arrest the heart in diastole) as described previously [7]. This was followed by perfusion fixation with freshly prepared fixative containing 2.5% glutaraldehyde, 2% paraformaldehyde in 0.15 M cacodylate buffer, and postfixed in 1% $OsO_4$ in 0.1 M cacodylate buffer for 1 hour on ice. After perfusion, small pieces of the left ventricle (in-between the apex and the middle of the heart) were immersed in the above fixative for 12-16 hrs. The tissues were stained *en bloc* with 2-3% uranyl acetate for 1 hour on ice. The tissues were dehydrated in graded series of ethanol (20-100%) on ice followed by one wash with 100% ethanol and two washes with acetone (15 min each) and embedding with Durcupan. Sections were cut at 50 to 60 nm on a Leica UCT ultramicrotome, and picked up on Formvar and carbon-coated copper grids. Sections were stained with 2% uranyl acetate for 5 minutes and Sato's lead stain for 1 minute. Grids were viewed using a JEOL JEM1400-plus TEM (JEOL, Peabody, MA) and photographed using a Gatan OneView digital camera with 4k x 4k resolution (Gatan, Pleasanton, CA).

For morphometric analyses, samples were blinded, and two people did measurements randomly from different sections as described previously [8]. The line segment tool in ImageJ was used to measure diameters of dense core vesicles (DCV) and the dense core (DC) inside the DCV. The free-hand tool was also used to measure areas by manually tracing around the DCV membrane and outer boundary of DC.

**Flow cytometry analysis.** Mouse heart perfusion was performed as previously described [9]. Briefly, after extracting the heart from the mouse thorax, coronary retrograde perfusion was used to efficiently digest the extracellular matrix with collagenase and protease XIV.

The ventricles were then isolated, mechanically dissociated and filtered into a single cell suspension. Cardiomyocytes and stromal cells were separated by gravity sedimentation. Cardiac stromal cells were stained with fluorescence-tagged antibodies to detect macrophages ($CD11b^{+}F4/80^{+}$). Data were analyzed using FlowJo software.

**Tissue macrophage depletion study.** To deplete tissue macrophages, mice were given clodronate liposomes (50 mg/kg body weight; catalog F70101C-N, FormuMax, Sunnyvale, CA) every 3 days via intraperitoneal injection. Mice treated with control liposomes were used as control.

**Bone marrow transplantation.** To generate irradiated chimeras, 10-12 week old WT or CST-KO NCD recipient mice received a lethal dose of 10 Gy radiation, followed by tail vein injection of 2 x $10^6$ bone marrow cells from either WT or CST-KO NCD donor mice as described previously [10]. After 16 weeks of bone marrow transplantation, all mice were subjected to non-invasive tail-cuff measurement of blood pressure.

**Real Time PCR.** Total RNA from heart tissue was isolated using RNeasy Mini Kit and reverse-transcribed using qScript cDNA synthesis kit. cDNA samples were amplified using PERFECTA SYBR FASTMIX L-ROX 1250 and analyzed on an Applied Biosystems 7500 Fast Real-Time PCR system. All PCRs were normalized to *Rplp0*, and relative expression levels were determined by the $\Delta\Delta C_t$ method.

**Langendorff perfused heart model.** Mice (mixed genetic background: 50% 129svJ/50% C57BL/6) were anesthetized with sodium pentobarbital (50 mg/kg intraperitoneal), the heart excised and the aorta cannulated for Langendorff perfusion of the coronary circulation, as described previously [11]. All hearts were perfused at a hydrostatic pressure of 80 mmHg with modified Krebs-Henseleit buffer bubbled with 95% $O_2$/5% $CO_2$ at 37°C, and containing: 119 mM NaCl, 22 mM $NaHCO_3$, 4.7 mM KCl, 2.5 mM $CaCl_2$, 1.2 mM $MgCl_2$, 1.2 mM $KH_2PO_4$, 11 mM D-glucose, and 0.5 mM EDTA. For the ischemia reperfusion (I-R) protocol a 15 min normoxic stabilization at intrinsic heart rates was followed by ventricular pacing at 7 Hz. After further 15 min hearts baseline measures were made before subjecting hearts to 25 min of global normothermic ischemia followed by 45 min aerobic reperfusion. For the ischemic preconditioning (IPC) protocol, the stabilization period was followed by two cycles of ischemia (5 min, no pacing) and reperfusion (5 min, pacing after 1 min re-perfusion).

**Measurement of catecholamines:** Mice were anesthetized by inhalation of isoflurane and blood was collected from the heart in potassium-EDTA tubes. Adrenal and plasma catecholamines were measured by ACQUITY UPLC H-Class System fitted with an Atlantis dC18 column (100A, 3 µm, 3 mm x 100 mm) and connected to an electrochemical detector (ECD model 2465) (Waters Corp, Milford, MA). The mobile phase (isocratic: 0.3 ml/min) consisted of phosphate-citrate buffer and acetonitrile at 95:5 (vol/vol). An internal standard 3.4-dihydroxybenzylamine (DHBA 400 ng) was added to the adrenal homogenate in 0.1N HCl. The ECD was set at 50 nA for determination of adrenal NE and EPI and at 2 nA for determination of adrenal DA. The ECD was set at 500 pA for determination of cardiac NE. For determination of plasma catecholamines, DHBA (2 ng) was added to 150 µl plasma and adsorbed with ~15 mg of activated aluminum oxide for 10 min in a rotating shaker. After washing with 1 ml water adsorbed catecholamines were eluted with 100 µl of 0.1N HCl. The ECD was set at 500 pA for determination of plasma catecholamines. The data were analyzed using Empower software (Waters Corp, Milford, MA). Catecholamine levels were normalized with the recovery of the internal standard. Catecholamines were expressed as nM (plasma) or nmol/mg protein (adrenal gland).

### SUPPLEMENTAL TABLE 1: PRIMER SEQUENCES

| **Gene** | **Gene ID** | **Forward primer (5'->3')** | **Reverse primer (5'->3')** |
|---|---|---|---|
| *Arg1* | 11846 | CTCCAAGCCAAAGTCCTTAGAG | GGAGCTGTCATTAGGGACATCA |
| *Ccl2* | 20296 | TTAAAAACCTGGATCGGAACCAA | GCATTAGCTTCAGATTTACGGGT |
| *Clec7a* | 56644 | GACTTCAGCACTCAAGACATCC | TTGTGTCGCCAAAATGCTAGG |
| *Clec10a* | 17312 | TGAGAAAGGCTTTAAGAACTGGG | GACCACCTGTAGTGATGTGGG |
| *Cxcl1* | 14825 | CTGGGATTCACCTCAAGAACATC | CAGGGTCAAGGCAAGCCTC |
| *Emr1* | 13733 | TGACTCACCTTGTGGTCCTAA | CTTCCCAGAATCCAGTCTTTCC |
| *Ifng* | 15978 | ATGAACGCTACACACTGCATC | CCATCCTTTTGCCAGTTCCTC |
| *Il4* | 16189 | GGTCTCAACCCCCAGCTAGT | GCCGATGATCTCTCTCAAGTGAT |
| *Il10* | 16153 | GCTCTTACTGACTGGCATGAG | CGCAGCTCTAGGAGCATGTG |
| *Il12b* | 16160 | TGGTTTGCCATCGTTTTGCTG | ACAGGTGAGGTTCACTGTTTCT |
| *Itgam* | 16409 | CCATGACCTTCCAAGAGAATGC | ACCGGCTTGTGCTGTAGTC |
| *Itgax* | 16411 | CTGGATAGCCTTTCTTCTGCTG | GCACACTGTGTCCGAACTCA |
| *Mrc1* | 17533 | CTCTGTTCAGCTATTGGACGC | CGGAATTTCTGGGATTCAGCTTC |
| *Nos2* | 18126 | GTTCTCAGCCCAACAATACAAGA | GTGGACGGGTCGATGTCAC |
| *Tnf* | 21926 | CCCTCACACTCAGATCATCTTCT | GCTACGACGTGGGCTACAG |

### SUPPLEMENTAL FIGURES

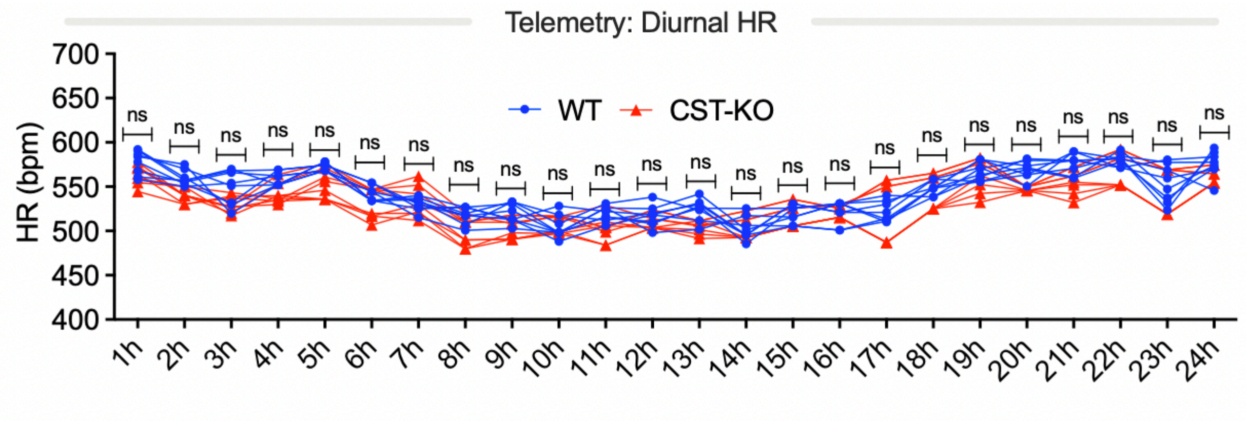


**Figure S1**

**Figure S1. Heart rate unaltered in CST-KO mice.** Diurnal heart rate (HR) by telemetry in wild-type (WT) and CST-KO mice (n=8). Two-way ANOVA followed by multiple comparison test: ns, not significant.

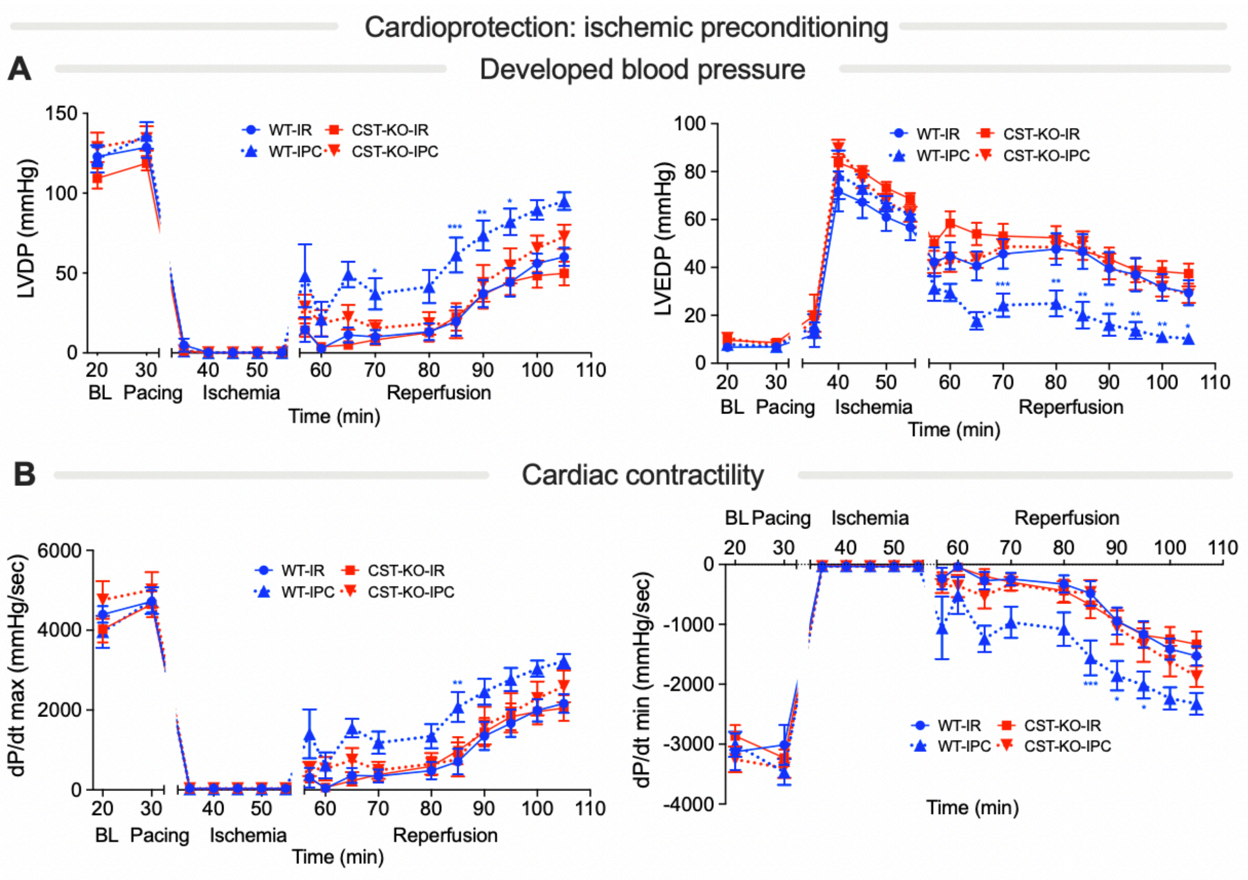

Cardioprotection: ischemic preconditioning
A
Developed blood pressure
LVDP (mmHg)
WT-IR
CST-KO-IR
WT-IPC
CST-KO-IPC
BL
Pacing
Ischemia
Reperfusion
Time (min)
LVEDP (mmHg)
B
Cardiac contractility
dP/dt max (mmHg/sec)
dP/dt min (mmHg/sec)


**Figure S2**

**Figure S2. Reduced ischemic preconditioning (IPC)-induced cardioprotection in CST-KO mice.** Hearts were excised and perfused on a Langendorff apparatus for ischemia/reperfusion (IR) and IPC studies (n=8-10). **(A)** Left ventricular developed pressure (LVDB) and left ventricular end diastolic pressure (LVEDP). **(B)** Maximum and minimum rates of pressure development in the left ventricle ($dP/dt_{max}$ and $dP/dt_{min}$). Two-way ANOVA followed by Tukey's multiple comparison test: *$p<0.05$; **$p<0.01$; ***$p<0.001$.

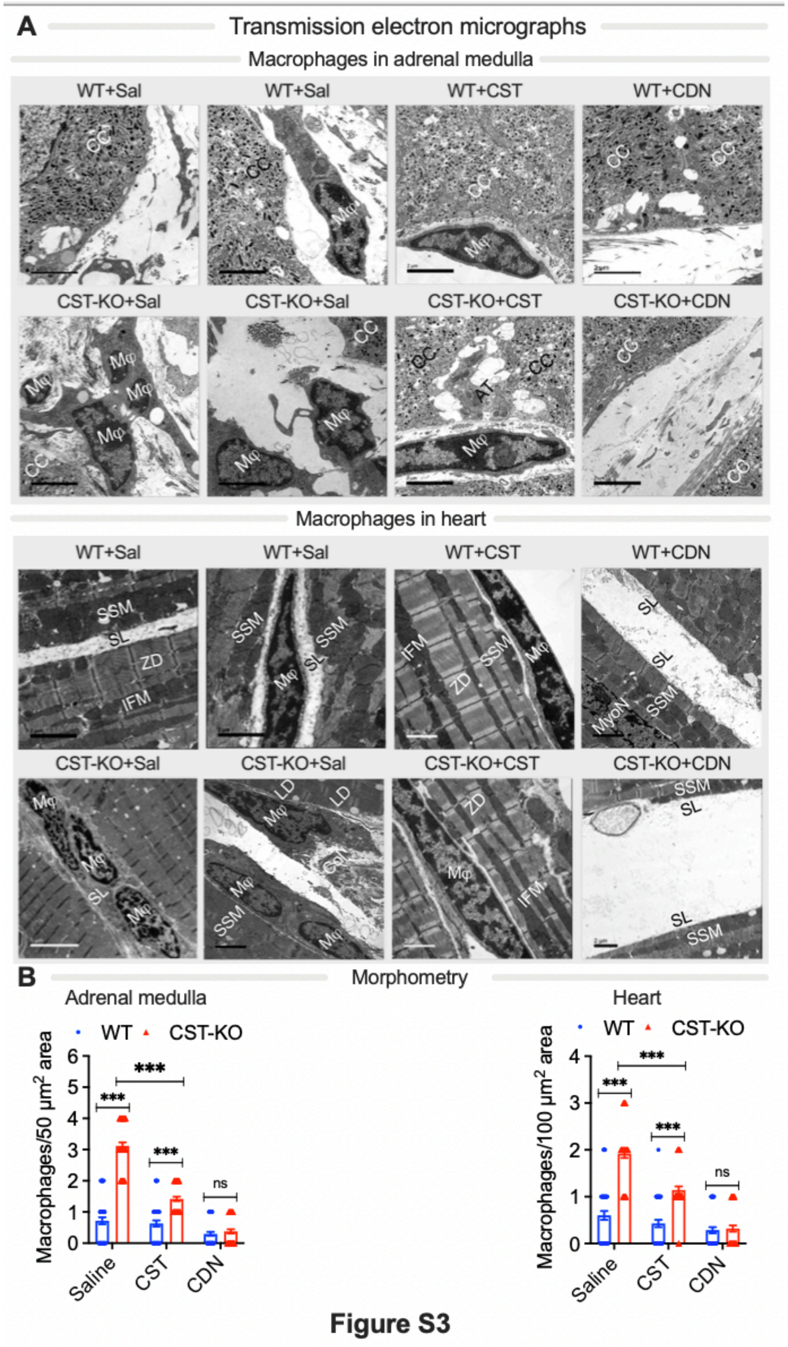
A
Transmission electron micrographs
Macrophages in adrenal medulla
WT+Sal
WT+Sal
WT+CST
WT+CDN
CST-KO+Sal
CST-KO+Sal
CST-KO+CST
CST-KO+CDN
Macrophages in heart
WT+Sal
WT+Sal
WT+CST
WT+CDN
CST-KO+Sal
CST-KO+Sal
CST-KO+CST
CST-KO+CDN
B
Morphometry
Adrenal medulla
WT
CST-KO
Macrophages/50 µm² area
Saline
CST
CDN
***
ns
Heart
Macrophages/100 µm² area

**Figure S3**

**Figure S3. Macrophage infiltration in adrenal gland and heart of CST-KO mice. (A)** Representative transmission electron microscopy (TEM) micrographs of the adrenal medulla and heart showing macrophages (MΦ) in WT and CST-KO mice. **(B)** Quantification of panel A (n=4). CDN: chlodronate. CST: intraperitoneal injection of CST. Sal: saline control. AT, axon terminal; CC, chromaffin cell; Col, collagen; IFM, intermyofibrillar mitochondria; MyoN, myocyte nucleus; Mϕ, macrophage; SL, sarcolemma; SSM, sub-sarcolemmal mitochondria; ZD, Z disc. Two-way ANOVA followed by Tukey's multiple comparison test: ns, not significant; ***$p<0.001$.

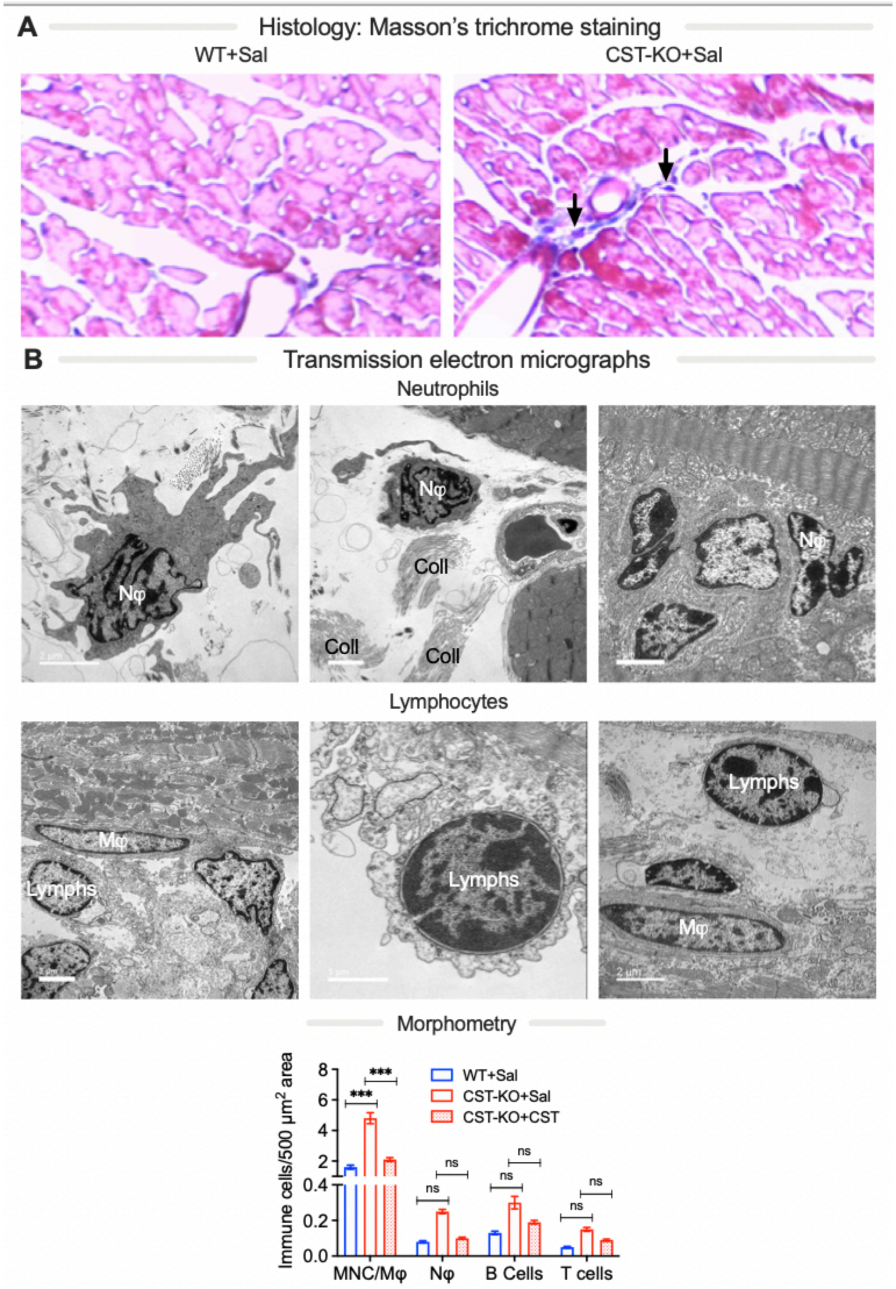
A
Histology: Masson's trichrome staining
WT+Sal
CST-KO+Sal
B
Transmission electron micrographs
Neutrophils
Nφ
Nφ
Coll
Coll
Coll
Nφ
Lymphocytes
Mφ
Lymphs
Lymphs
Lymphs
Mφ
Morphometry
Immune cells/500 µm2 area
WT+Sal
CST-KO+Sal
CST-KO+CST
***
***
ns
MNC/Mφ
Nφ
B Cells
T cells

**Figure S4**

**Figure S4. Increased macrophage infiltration in heart of CST-KO mice. (A).** Masson's trichrome stained micrographs of the left ventricle showing infiltration of immune cells (shown by arrows) in CST-KO mice. WT: wild-type mice; Sal, saline control. **(B)** Representative transmission electron microscopy (TEM) micrographs showing neutrophils and lymphocytes. **(C)** Quantification of panel B. Coll, collagen; MNC, mononuclear cells; Lymphs, lymphocytes; M$\varphi$, macrophages; N$\varphi$, neutrophils. Two-way ANOVA followed by Tukey's multiple comparison test: ns, not significant; ***p<0.001.

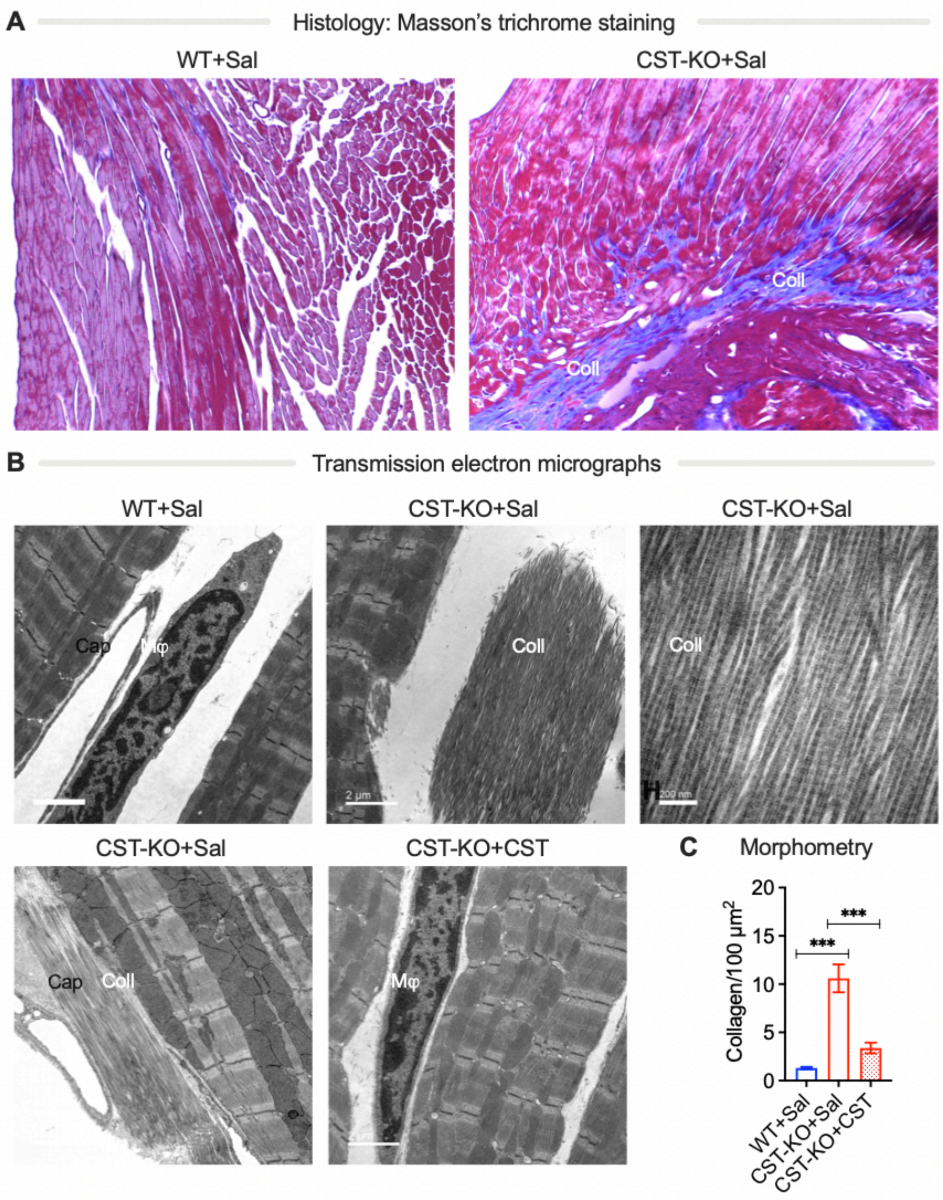
A
Histology: Masson's trichrome staining
WT+Sal
CST-KO+Sal
Coll
Coll
B
Transmission electron micrographs
WT+Sal
CST-KO+Sal
CST-KO+Sal
Cap
Mφ
Coll
Coll
2 μm
200 nm
CST-KO+Sal
CST-KO+CST
Cap
Coll
Mφ
C
Morphometry
Collagen/100 μm²
20
15
10
5
0
***
***
WT+Sal
CST-KO+Sal
CST-KO+CST

**Figure S5**

**Figure S5. Increased fibrosis in heart of CST-KO mice. (A).** Masson's trichrome stained micrographs showing fibrosis in the left ventricle of the CST-KO mice. CST-KO and wild-type (WT) mice were treated with saline (Sal) or CST. **(B)** Representative transmission electron microscopy (TEM) micrographs showing the presence of collagen fibers (fibrosis) in CST-KO heart. **(C)** Quantification of panel B. Cap, capillary; Coll, collagen; M$\varphi$, macrophages. One-way ANOVA followed by multiple comparison test: ***$p<0.001$.

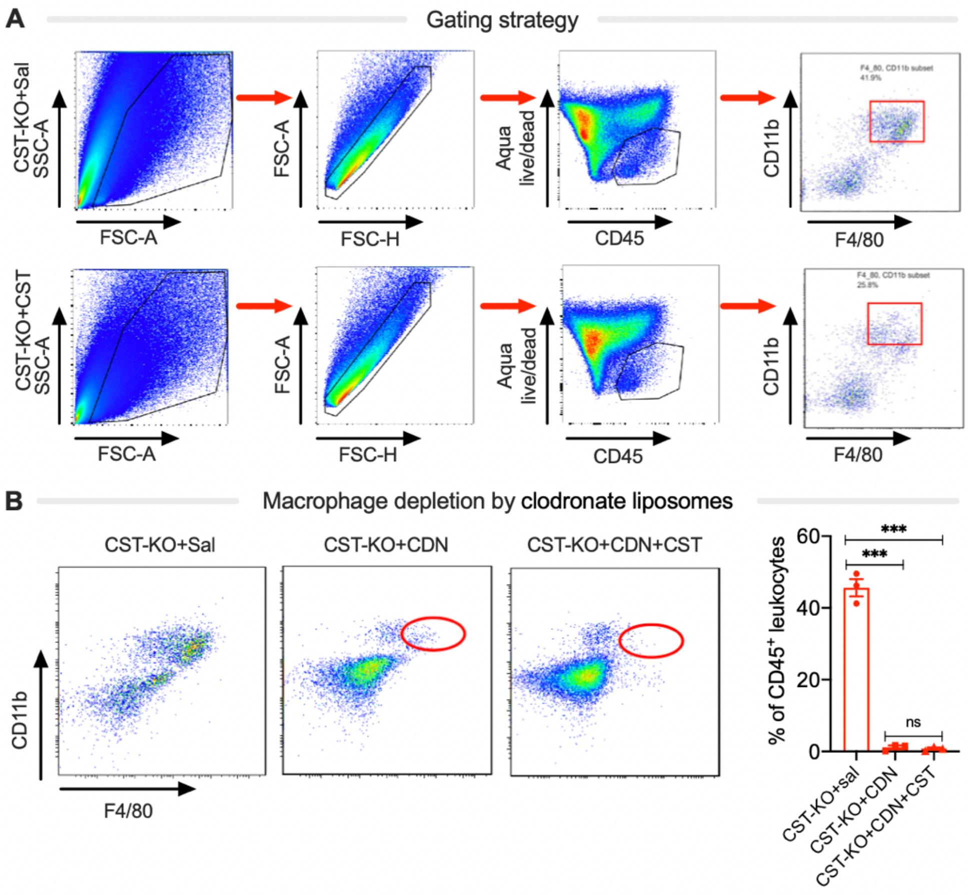
A
Gating strategy
CST-KO+Sal
SSC-A
FSC-A
FSC-A
FSC-H
Aqua live/dead
CD45
CD11b
F4/80
CST-KO+CST
B
Macrophage depletion by clodronate liposomes
CST-KO+Sal
CST-KO+CDN
CST-KO+CDN+CST
% of CD45+ leukocytes
***
ns
CST-KO+sal
CST-KO+CDN
CST-KO+CDN+CST

**Figure S6**

**Figure S6. Macrophage infiltration in heart of CST-KO mice by flow cytometry. (A)** Gating strategy for flow cytometric analysis of cardiac macrophages. **(B)** Depletion of macrophages by clodronate liposomes (CDN). Mice were treated with saline (Sal) or CST. The cardiac macrophage population was evaluated by flow cytometry analysis after treatment of clodronate liposomes (n=4 mice per group). Cardiac macrophage: $CD45^+CD11b^+F4/80^+$. SSC-A, side scatter area; FSC-A, forward scatter area; FSC-H, forward scatter height. One-way ANOVA followed by multiple comparison test: ns, not significant; ***$p<0.001$.

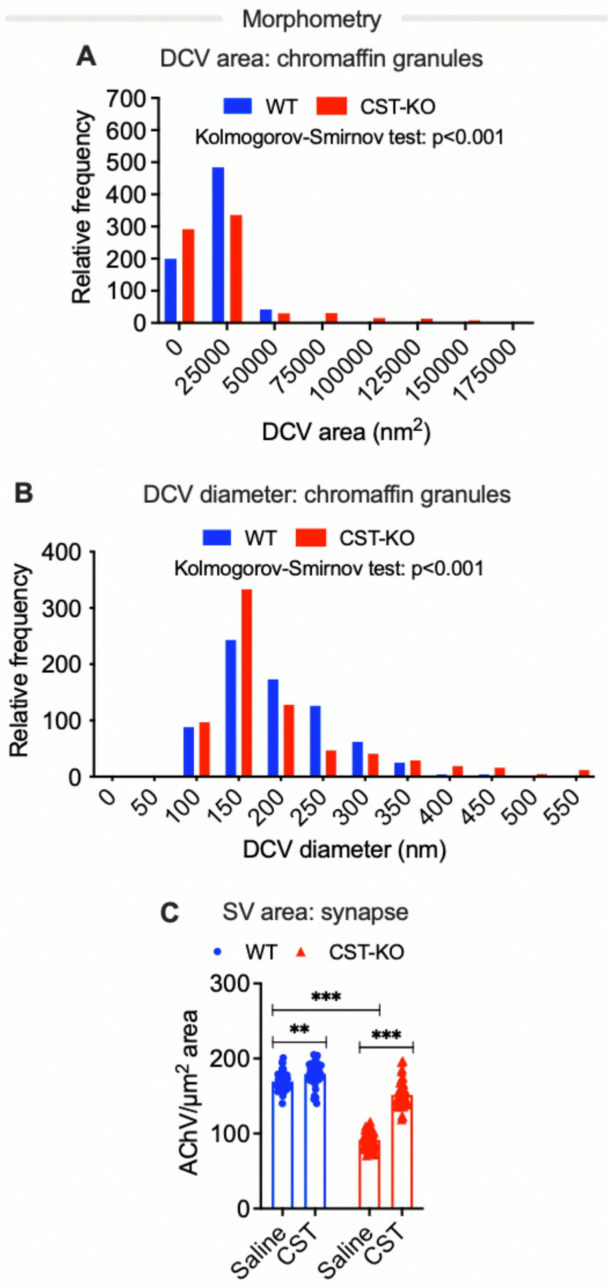
Morphometry
A
DCV area: chromaffin granules
WT
CST-KO
Kolmogorov-Smirnov test: p<0.001
Relative frequency
700
600
500
400
300
200
100
0
0
25000
50000
75000
100000
125000
150000
175000
DCV area (nm2)
B
DCV diameter: chromaffin granules
WT
CST-KO
Kolmogorov-Smirnov test: p<0.001
Relative frequency
400
300
200
100
0
0
50
100
150
200
250
300
350
400
450
500
550
DCV diameter (nm)
C
SV area: synapse
WT
CST-KO
AChV/μm2 area
300
200
100
0
***
**
***
Saline
CST
Saline
CST

**Figure S7**

**Figure S7. Differential vesicle area and diameter in CST-KO mice. (A-B)** Quantification of main figure 6B with areas **(A)** and diameters **(B)** of dense core vesicles (DCV; n=728). **(C)** Quantification of main figure 6C with density of AChVs (n=50). Kolmogorov-Smirnov test (A&B), and two-way ANOVA (C): **p<0.01; ***p<0.001.

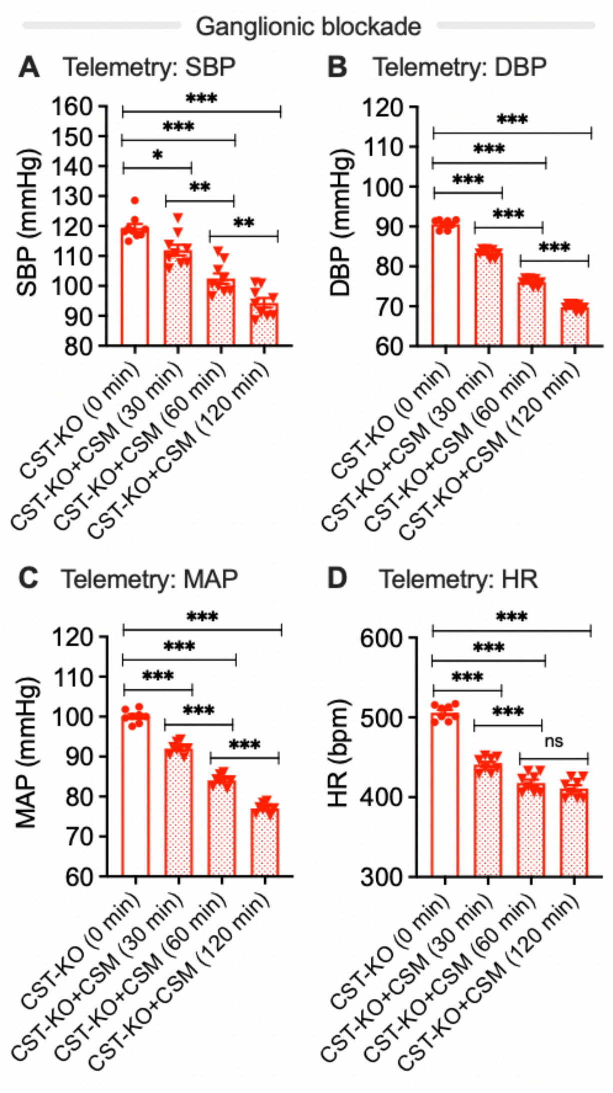
Ganglionic blockade
A Telemetry: SBP
B Telemetry: DBP
C Telemetry: MAP
D Telemetry: HR
SBP (mmHg)
DBP (mmHg)
MAP (mmHg)
HR (bpm)
CST-KO (0 min)
CST-KO+CSM (30 min)
CST-KO+CSM (60 min)
CST-KO+CSM (120 min)
ns

**Figure S8**

**Figure S8. Reversal of elevated blood pressure in CST-KO mice by the nicotinic acetylcholine receptor antagonist chlorisondamine.** (**A**) Systolic blood pressure (SBP), (**B**) diastolic blood pressure (DBP), (**C**) mean arterial pressure (MAP), and (**D**) heart rate (HR) after 30, 60, and 120 min of chlorisondamine treatment (5 μg/g body weight, intraperitoneal) in telemetered CST-KO mice. One-way ANOVA: *$p<0.05$; **$p<0.01$; ***$p<0.001$.

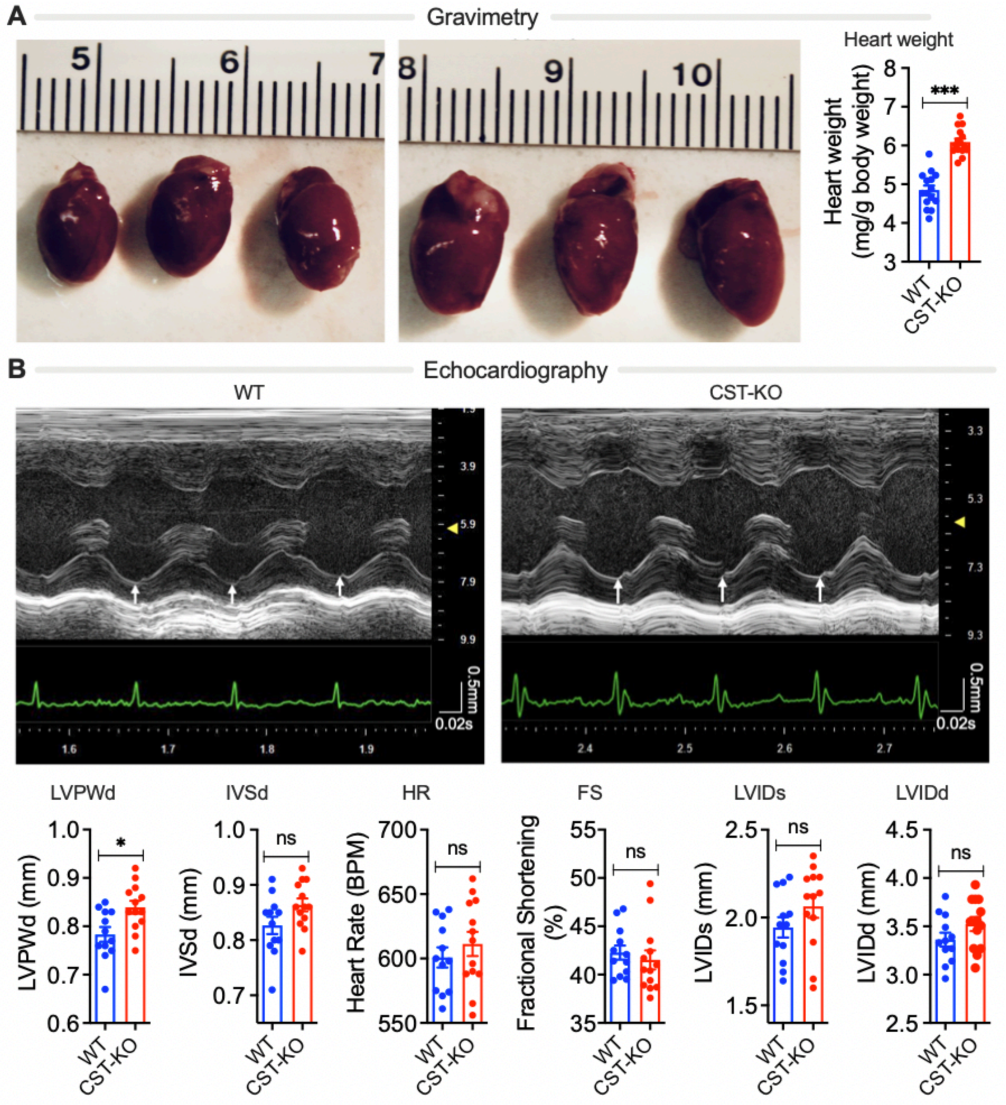
A
Gravimetry
Heart weight
Heart weight (mg/g body weight)
***
WT
CST-KO
B
Echocardiography
WT
CST-KO
0.5mm
0.02s
LVPWd
IVSd
HR
FS
LVIDs
LVIDd
LVPWd (mm)
IVSd (mm)
Heart Rate (BPM)
Fractional Shortening (%)
LVIDs (mm)
LVIDd (mm)
*
ns

**Figure S9**

**Figure S9. Left ventricular hypertrophy in CST-KO mice. (A)** Representative images and weights of wet hearts (n=16). **(B)** Representative echocardiograms from the left ventricle (LV) of age-matched male wild-type (WT; n=12) and CST-KO (n=13) mice showing increased posterior wall thickness (shown by white arrows) in CST-KO mice. LV posterior wall thickness at end-diastole (LVPWd), interventricular septum thickness at end-diastole (IVSd), heart rate (HR), fractional shortening (FS), LV internal diameter during systole (LVIDs), and LV internal diameter during diastole (LVIDd). Unpaired *t*-test: ns, not significant; *$p<0.05$; **$p<0.01$; ***$p<0.001$.
